# An animal model for autoinflammation with infantile enterocolitis

**DOI:** 10.1101/2024.11.16.623944

**Authors:** Yuhang Wang, Joyce Z. Gao, Prajwal Gurung, Sarah P. Short, Yiqin Xiong, Zizhen Kang

## Abstract

Inflammasomes, particularly NLRC4, play a crucial role in immune responses to intracellular bacterial infections. However, gain-of-function mutations of NLRC4 are linked to severe autoinflammatory diseases, including autoinflammation with infantile enterocolitis (AIFEC). Existing mouse models do not adequately replicate the chronic, unprovoked inflammation observed in AIFEC patients. In this study, we developed a loxP-flanked NLRC4 V341A knock-in (KI) mouse model. Our global NLRC4 V341A KI mice exhibited symptoms closely mirroring those of human AIFEC. These mice demonstrated severe infantile enterocolitis, characterized by heightened intestinal inflammation, compromised intestinal barrier integrity, disrupted gut epithelium, and severe diarrhea, with high mortality within 10 days postnatally. Additionally, they displayed systemic autoinflammation marked by elevated levels of IL-1β, IL-18, and IL-6, alongside cytopenia and hemophagocytosis. In contrast, adult NLRC4 conditional KI mice exhibited autoinflammation with only mild enterocolitis. Since AIFEC is characterized by life-threatening enterocolitis in infancy and recurrent severe autoinflammation throughout life, our conditional NLRC4 KI model effectively recapitulates the clinical features of human AIFEC. Moreover, the NLRC4 V341A-mediated monogenic infantile enterocolitis observed in our model resembles infantile-onset inflammatory bowel disease (IBD), positioning it as a valuable platform for research on very early-onset IBD.

## Introduction

The NLRC4 inflammasome is a crucial multiprotein complex involved in the immune response, particularly in detecting specific pathogens such as *Salmonella typhimurium*. As part of the larger inflammasome family, it plays a significant role in activating inflammatory processes^1^. The NAIP-NLRC4 inflammasome specifically responds to cytosolic flagellin, as well as the needle proteins and inner rod proteins of the bacterial type III secretion system. Upon activation, the NLRC4 inflammasome assembles and triggers a robust inflammatory response characterized by the production of IL-1β and IL-18, along with pyroptosis—a highly inflammatory form of programmed cell death^2^. Pyroptosis, initiated by cleaved gasdermin D (GSDMD), creates membrane pores that eliminate infected cells and inhibit pathogen replication^3, 4^. Moreover, caspase-1 cleaves pro-cytokines such as pro-IL-1β and pro-IL-18 into their active forms, which are essential for inducing fever, recruiting immune cells, and promoting tissue inflammation. NLRC4 is expressed in various cell types, with particularly high expression in myeloid and epithelial cells^5, 6^.

While the NLRC4 inflammasome is vital for immune defense, excessive activation can result in pathological inflammation, leading to NLRC4-associated autoinflammatory disorders or NLRC4 inflammasomopathies^5^. Two inborn errors of NLRC4 were reported simultaneously in 2014, one variant is NLRC4 T337S^7^ and the other is NLRC4 V341A^8^. These mutations were linked to a condition later termed autoinflammation with infantile enterocolitis (AIFEC). Since then, fourteen pathogenic NLRC4 variants have been reported. Clinical phenotypes have expanded from AIFEC to familial cold autoinflammatory syndrome (FCAS4) and neonatal-onset multisystem inflammatory disease (NOMID). Among them, AIFEC is the most aggressive with lethal consequences^9-13^. AIFEC patients typically exhibit signs of autoinflammation, including elevated ferritin levels, pancytopenia, and hemophagocytosis, particularly manifesting as macrophage activation syndrome (MAS) with markedly elevated peripheral IL-18^7, 8^. Another hallmark of AIFEC is the development of infantile enterocolitis, characterized by diarrhea, abdominal pain, villus blunting, and mixed inflammatory infiltrates which could be throughout the intestine^8^. Notably, all reported cases of NLRC4-mediated enterocolitis occurred in infancy^7, 8, 14, 15^, with the colitis in surviving patients often resolving or becoming milder. In contrast, recurrent autoinflammation persists into adulthood^13^. High mortality rates have been observed among infants with AIFEC, yet standardized therapies remain lacking^12, 13^. Several gain-of-function NLRC4 variants can contribute to AIFEC, with the NLRC4 V341A mutation being particularly prevalent. This mutation occurs at amino acid position 341 within the autoinhibitory HD-1 subdomain of the NOD of NLRC4. Crystal structure analyses suggest that hydrophobic residues at this position are critical for closing the ‘lid’ over the ADP binding pocket, thereby preventing ADP–ATP exchange. Consequently, the V341A mutation may disrupt this inhibitory conformation, leading to constitutive activation of the NLRC4 inflammasome ^8^.

To elucidate the mechanisms underlying AIFEC and develop targeted treatments to mitigate hyperinflammatory responses, a mouse model that accurately recapitulates the disease’s symptoms is critically needed. However, current NLRC4 models fail to replicate the spontaneous, chronic inflammation and neonatal-onset enterocolitis observed in patients with NLRC4 V341A gain-of-function mutations, hindering progress in understanding and treating the disease. In this study, we developed a loxP-flanked NLRC4 V341A knock-in (KI) mouse model, incorporating the V341A (GTG to GCG) mutation into the mouse NLRC4 gene. The global NLRC4 KI mice exhibited severe infantile enterocolitis and systemic autoinflammation, closely mirroring the symptoms observed in human AIFEC. Remarkably, most KI pups died within 10 days postnatally, showing profound intestinal inflammation, diarrhea, multi-organ damage, and hypoglycemia. Hyperactivation of the NLRC4 inflammasome was detected in both the gut epithelium and macrophages of KI mice, accompanied by significantly elevated serum levels of IL-1β, IL-18, and IL-6, as well as cytopenia and hemophagocytosis. Additionally, inducible expression of the NLRC4 V341A mutation in adult mice led to autoinflammation, though milder colitis. Importantly, we found that inhibiting LRRK2 kinase activity—an upstream regulator of NLRC4 inflammasome activation—provides a promising strategy to suppress hyperactivation of the NLRC4 inflammasome. This study introduces a valuable mouse model that recapitulates key features of AIFEC, paving the way for new investigations into the disease’s pathogenesis and revealing potential therapeutic targets.

## Materials and Methods

### Animals

The NLRC4 V341A-Flag^flox/flox^ mice was generated by Cyagen Biosciences using Cyagen’s TurboKnockout^®^ gene targeting service through homologous recombination. The Nlrc4 gene (NCBI Reference Sequence: NM_001033367.3) is located on mouse chromosome 17. Nine exons have been identified, with the ATG start codon in exon 2 and the TAA stop codon in exon 9. In the targeting vector, the coding sequence of exon 2 was replaced with the “loxP-stop-loxPkozak-mutant Nlrc4 CDS-3*Flag-IRES-EGFP-polyA”. The V341A (GTG to GCG) mutation was introduced into mutant Nlrc4 CDS. In the targeting vector, the Neo cassette was flanked by SDA (self-deletion anchor) sites. DTA (Diphtheria toxin A) was used for negative selection. C57BL/6 ES cells were used for gene targeting. The final targeting vector was sequenced for validation. The NLRC4 V341A-Flag^flox/flox^ mice were genotyped by PCR and further confirmed by sequencing. After breeding with Cre transgenic mice in our facility, western blot was conducted to prove the expression at protein level by using Flag Tag antibody and NLRC4 antibody as shown in Figure 1. E2a-Cre (JAX, #003724)^16^ and R26-CreERT2 mice (JAX,008463)^16^ were purchased from the Jackson Laboratory. Both male and female mice were used. Littermate controls were used for all the experiments. The mouse ages are indicated in figure legends. All mice were bred and maintained in individually ventilated cages under specific pathogen-free conditions in accredited animal facilities. Animal experiments were approved by the Institutional Animal Care and Use Committee of the University of Iowa.

**Figure 1.**
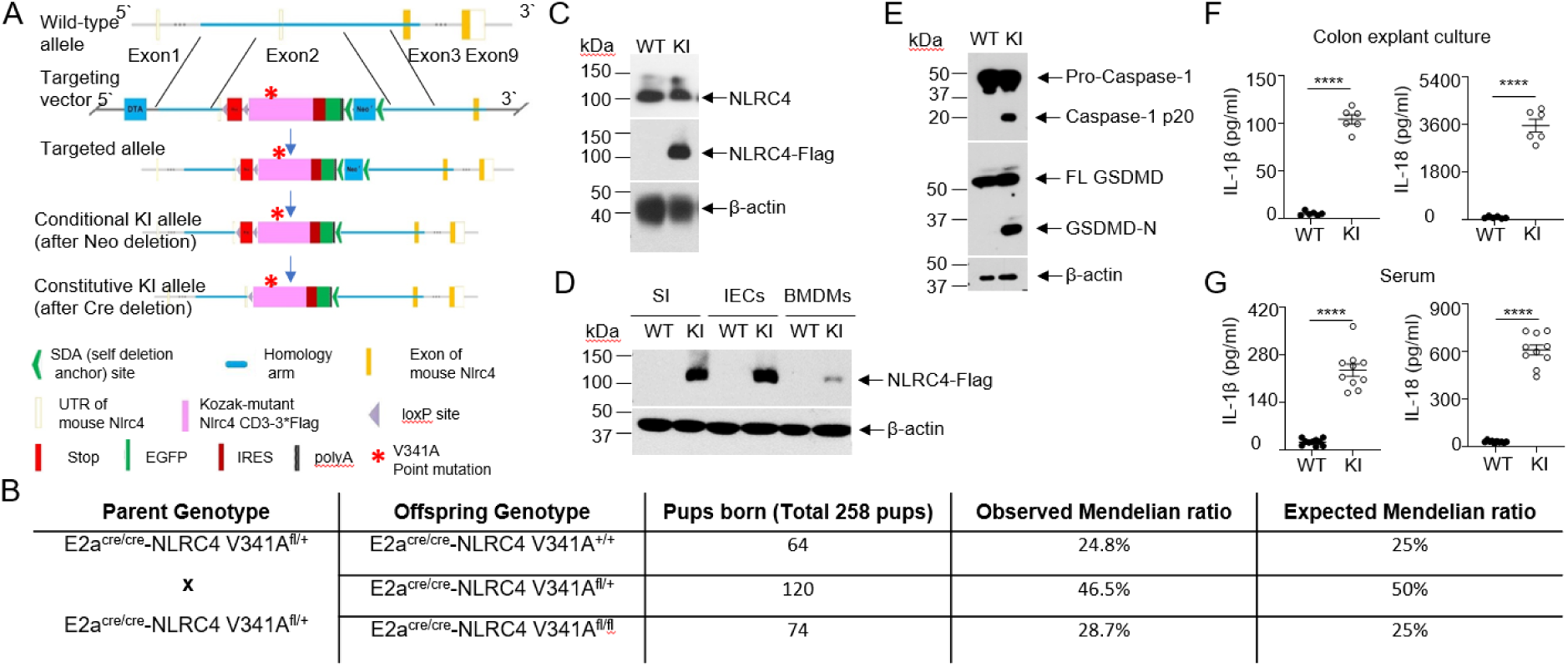
NLRC4 V341A KI mice manifest inflammasome hyperactivation. (A) Diagram illustrating the design and generation of NLRC4 V341A-Flag knock-in (KI) mice. (B) Mendelian inheritance analysis of NLRC4 KI offsprings as indicated. (C) Genotyping colon tissues of 6 days old E2aCre-NLRC4 V341A^fl/fl^ (KI) and E2aCre-NLRC4^+/+^ (wild-type, WT) mice by western blot. (D) Western blot analysis of NLRC4 V341A-Flag expression in indicated tissues, probed by Flag and β-actin antibodies. SI: Small intestine, IECs: Intestinal epithelial cells, and BMDMs: Bone marrow-derived macrophages. (E) Inflammasome activation in IECs from NLRC4 WT and KI mice was analyzed by western blot with indicated antibodies (FL GSDMD, full length Gasdermin-D; GSDMD-N, N-terminal GSDMD. (F) IL-1β and IL-18 levels in colon explant cultures from NLRC4 WT and KI mice were measured by ELISA (n=6 per group). Colon tissues were cultured in 1 ml of medium overnight, and the supernatant was collected for analysis. (G) IL-1 β and IL-18 levels of NLRC4 WT and KI mouse serum were measured by ELISA (n=10 per group). In panel F-G, data are shown as mean ± SEM; ****, p<0.0001 were determined by Student’s t test. Data are representative of at least three independent experiments.

### Culture of Bone marrow-derived macrophages (BMDMs)

Bone marrow was harvested from the femurs of mice by flushing with RPMI-1640 medium containing 2% FBS. The harvested cells were centrifuged at 300 × g for 10 minutes, followed by treatment with ACK lysis buffer (ThermoFisher, A1049201) for 30 seconds to lyse red blood cells. To neutralize the lysis buffer, 10 volumes of PBS were added, and the cells were centrifuged again at 300 × g for 10 minutes. The resulting cell pellet was resuspended in BMDM culture medium (RPMI-1640 supplemented with 10% FBS, antibiotics, L-glutamine, and 10 ng/ml M-CSF) and plated at a density of 1 × 10^6^ cells/ml in tissue culture plates. The culture medium was refreshed on days 4 and 6, and the BMDMs were used on day 7.

### Colonic explant culture

The whole colons were harvested, thoroughly rinsed with serum-free DMEM medium, and weighed to determine their initial weight. The collected colon tissues were cut into 2 mm pieces and then cultured as explants in regular RPMI 1640 medium supplemented with 10% FBS, L-glutamine, penicillin, and streptomycin, and placed in a standard cell-culture incubator for 24 hours. After the culture, the cell-free supernatants were obtained by centrifuging at 12,000 × g for 10 minutes at 4°C and stored in aliquots at −20°C for further analysis.

### Blood Chemical parameters assay

Chemical parameters of blood from NLRC4 WT and KI mice were measured as indicated (ALT, alanine aminotransferase; BUN, blood urea nitrogen; LDH, Lactate Dehydrogenase etc) by service from IDEXX Inc using VetScan VS2 analyzer.

### ELISA analysis

Serum levels of ferritin, AST, hemoglobin, as well as IL-6, IL-1β, and IL-18 in both serum and colon explant culture, were quantified using commercial ELISA kits according to the manufacturer’s instructions. The kits used were Ferritin (Catalog #80636, Crystal Chem), AST (Catalog #XPEM0857, XpressBio), Hemoglobin (Catalog #80640, Crystal Chem), IL-1β (Catalog #DY401-050, R&D), IL-6 (Catalog #DY406-05, R&D), and IL-18 (Catalog #7625-05, R&D). Cytokine levels in colon explant supernatants were normalized to the weight of the colon tissue.

### Peripheral blood cell count using hemocytometer

Peripheral blood samples were collected from NLRC4 WT and KI mice into EDTA-coated tubes to prevent coagulation. Peripheral blood cell counts were performed using a hemocytometer, following established methods from previous research^17^.

#### Red Blood Cell (RBC) Counting

Blood samples were diluted using an RBC diluting solution (Hayem ‘s Solution) consisting of mercuric chloride (0.5 g), sodium chloride (1 g), sodium sulfate (5 g), and distilled water (200 mL). A clean RBC pipette was used to draw blood to mark 0.5, followed by filling the pipette to mark 101 with the RBC diluting solution, creating a 1:200 dilution. The mixture was thoroughly mixed, and the first few drops were discarded. The diluted sample was then loaded into the hemocytometer, and the cells were allowed to settle. RBCs were counted in the central large square under a microscope at 40x magnification. The total RBC concentration was calculated using the standard hemocytometer formula, accounting for dilution, area, and chamber depth.

#### Platelet Counting

A platelet diluting solution of 1% ammonium oxalate was prepared by dissolving 1 g of ammonium oxalate in 100 mL of distilled water. The RBC pipette was pre-cleaned with alcohol before drawing blood to mark 0.5, and the pipette was then filled to mark 101 with the platelet diluting solution, creating a 1:200 dilution. After thoroughly mixing the sample, the first few drops were discarded, and the diluted blood was loaded into the hemocytometer. Platelets were counted in the central large square under a microscope at 10x magnification, and the total platelet concentration was calculated using the standard hemocytometer formula.

#### White Blood Cell (WBC) Counting

WBC counting was performed using a hemocytometer, a WBC pipette, and a WBC diluting solution composed of glacial acetic acid (1.5 mL), crystal violet solution (1% aqueous, 1 mL), and distilled water (98 mL). The WBC pipette was cleaned with alcohol, and blood was drawn up to mark 0.5. After removing any excess blood from the outside of the pipette, the pipette was filled to mark 11 with the WBC diluting solution, resulting in a 1:20 dilution. The mixture was then thoroughly mixed, and the first drops were discarded before loading the diluted sample into the hemocytometer. WBCs were counted in all four corner large squares under a microscope at 10x magnification, and the total WBC concentration was calculated using the standard hemocytometer formula.

### Isolation of intestinal epithelial cells (IECs)

intestinal epithelial cells (IECs) were isolated following a standard method as previously described^18^. The large intestine was carefully removed, and any attached tissue was trimmed away. The intestine, excluding the cecum, was opened lengthwise and cut into 1 cm pieces, which were then rinsed three times with ice-cold PBS. These pieces were further cut into 5 mm segments and placed in a solution of 5 mM EDTA/PBS at 4°C on a rocking platform for 30 minutes. The crypts were then released by shaking the tubes for 2 minutes and collected by spinning at 200 × g for 10 minutes at 4°C.

### Isolation of macrophages from spleen

Macrophages were isolated from mouse spleen using a standardized protocol to ensure high purity and viability. Spleens were carefully harvested from mice and placed in cold PBS. The spleen tissue was homogenized by gently pressing it through a 70 μm cell strainer using the plunger of a syringe. The cell suspension was then centrifuged at 300 x g for 5 minutes at 4°C to pellet the cells. Red blood cells were lysed using ACK lysis buffer for 5 minutes at room temperature, followed by another round of centrifugation and washing with cold PBS. After washing, the cell suspension was enriched for macrophages by using magnetic cell separation (MACS) technology. F4/80^+^ macrophages were positively selected using Microbeads (Catalog# 130-110-443, Miltenyi Biotec) according to the manufacturer’s instructions. The purified macrophages were then lysed in TRIzol reagent (Invitrogen) for RNA extraction and Q-PCR.

### Isolation of lamina propria cells

Lamina propria cells were isolated using a modified version of a previously described protocol^19^. Briefly, large intestines, cleared of Peyer’s patches, were gently removed from the abdominal cavity. The mesentery and associated fatty tissue were carefully excised using forceps. Intestines were then opened longitudinally and cut into 1 cm segments. These segments were thoroughly washed with room temperature PBS three times. After washing, the intestinal pieces were incubated in a solution of PBS plus 30 mM EDTA and10 mM HEPES at 37°C on a rocking platform for 10 minutes. The epithelial cell layer was removed by shaking the tissue for 2 minutes. The remaining intestinal fragments were further cut into 5 mm pieces and digested in a solution containing collagenase VIII (200 U/ml, Sigma-Aldrich) and DNase I (150 μg/ml, Sigma-Aldrich) in RPMI 1640 complete medium for 50 minutes. Following digestion, cells were resuspended in 4 ml of 40% Percoll (GE Healthcare) and layered onto 2.5 ml of 80% Percoll in a 15 ml tube. Cells were collected from the interphase of the Percoll gradient, washed twice, and resuspended in PBS with 1% FBS. The isolated lamina propria cells were then prepared for flow cytometry analysis.

### Colon diameter measurement

To measure colon diameter, distal colon sections were fixed, embedded in paraffin, sectioned, and examined under a microscope using calibrated imaging software to determine external diameters. Multiple measurements were obtained per section, and the average diameter was calculated and recorded.

### Feces wet/dry ratio

The feces wet/dry ratio was determined using a modified version of a previously described protocol^20^. Briefly, fecal samples were collected from the contents of the small intestine and colon. Each sample was initially weighed to obtain wet weight (WW), then dried in a 70°C incubator until a constant weight was achieved. The dried samples were subsequently weighed to determine the dry weight (DW). The wet/dry ratio was calculated by dividing the initial wet weight by the final dry weight (WW/DW).

### Flow cytometry

Isolated lamina propria cells were resuspended in PBS containing 5% FBS and stained with fluorescence-conjugated antibodies to CD4 (APC-cy7-CD4, Catalog#100526, clone RM4-5), CD8 (APC-CD8, Catalog#100712, clone 53-6.7, Bio-legend), CD45 (Percp-CD45, Catalog#103129, clone 30-F11), and Ly6G (APC-Ly6G, Catalog#127613, clone 1A8), F4/80 (PE-F4/80, Catalog#MCA497, clone Cl:A3-1, Bio-Rad) and isotype controls, purchased from BD Biosciences. Antibodies were diluted at 1:100 when used. Hemophagocytosis in the spleen was analyzed by flow cytometry. Splenocytes isolated from 6-day-old NLRC4 wild-type (WT) and knock-in (KI) mice were initially stained with fluorescence-conjugated antibodies targeting CD11b (percp-CD11b, BioLegend, Catalog#101230, clone M1/70). After surface staining, cells were incubated with unconjugated Ter119 antibody (Catalog#116202, BioLegend) to block the respective surface antigens. Cells were then permeabilized to allow intracellular staining for phagocytosed Ter119 (FITC-Ter119, Catalog#116205, BioLegend) cells. Flow cytometry was performed using a Cytek Aurora, and the data were analyzed with FlowJo software.

### Immunoblot

Colon tissues or BMDMs were lysed using radioimmunoprecipitation (RIPA) assay buffer, which was supplemented with Complete Mini Protease Inhibitor Cocktail and Phosphatase Inhibitor Cocktail from Roche. The lysates were put on ice for 30 minutes and vortexed every 5 minutes. Following centrifugation at 13,000 rpm for 10 minutes at 4°C, the supernatants were collected. Protein concentration was determined by a BCA Protein Assay Kit from Pierce. Subsequently, the proteins were resolved by SDS-PAGE and transferred to a 0.45-mm PVDF membrane. For immunoblot analysis, the specified primary antibodies were used at a 1:1000 dilution. Primary antibodies: anti-caspase-1 antibody (Catalog#AG-20B-0042-C100, AdipoGen); anti-GSDMD antibody (Catalog#ab209845, Abcam); anti-mouse IL-1β antibody (Catalog#AF-401-NA, R&D); anti-β-actin antibody (Catalog#3700S, CST); Anti-NLRC4 antibody (Catalog#061125, Sigma-Aldrich); Anti-Flag antibody (Catalog#F1804, Sigma-Aldrich); p-MLKL (Catalog#91689S, CST); MLKL (Catalog#26539T, CST); anti-caspase3 antibody (Catalog#9661, CST). Horseradish peroxidase (HRP)-conjugated secondary antibodies were selected according to the host species of the primary antibodies. The membranes were treated with ECL reagent (Catalog#34580, Thermo Fisher Scientific), and signal detection was performed by exposure to X-ray film (Z&Z Medical, Fuji X-Ray Film).

### Quantitative PCR

Colon tissues or isolated macrophages from the spleen were carefully collected and homogenized using TRIzol reagent (Invitrogen) to ensure efficient RNA extraction. RNA was isolated according to the manufacturer’s instruction. The high-quality RNA (500 ng) was then reverse transcribed into complementary DNA (cDNA) using the High-Capacity cDNA Reverse Transcription Kit (Applied Biosystems). Quantitative PCR (Q-PCR) was subsequently carried out using SYBR Green Real-time PCR Master Mix (Catalog#K1070, APExBio) on a Real-Time PCR System (Applied Biosystems). Each reaction was performed in triplicate in a 10 μl volume, containing 5 μl of SYBR Green mix, 0.3 μM of forward and reverse primers, and 0.5 μl of cDNA. The relative gene expression levels were calculated using the comparative Ct (ΔΔCt) method, normalized to the expression of a housekeeping gene β-actin.

### Immunohistochemistry

Formalin-fixed, paraffin-embedded distal small intestine tissue sections were carefully processed for immunohistochemistry (IHC). First, sections were deparaffinized by immersion in xylene (3 changes, 5 minutes each) followed by rehydration through a graded ethanol series (100%, 95%, 70%, and 50%) and finally rinsed in distilled water. To unmask antigens, antigen retrieval was performed by heating the sections in citrate antigen retrieval buffer (pH 6.0) at 95°C for 30 minutes in a pressure cooker, followed by gradual cooling to room temperature. After cooling, tissue sections were permeabilized with 0.1% Triton X-100 in PBS for 10 minutes, and non-specific binding sites were blocked with 5% normal goat serum in PBS for 1 hour at room temperature. The sections were then incubated overnight at 4°C with primary antibodies, including anti-lysozyme (Catalog #sc-518012, Santa Cruz Biotechnology, 1:500 dilution) and anti-ZO-1 (Catalog #14-9776-82, eBioscience,1:500 dilution). Control sections were incubated with isotype-matched IgG antibodies. The following day, the sections were washed three times with PBS and incubated for 1 hour at room temperature with fluorescence-conjugated secondary antibodies. After secondary antibody incubation, the sections were washed, counterstained with DAPI (4’,6-diamidino-2-phenylindole) to visualize nuclei and mounted using anti-fade mounting medium. Fluorescent images were captured using an Olympus DP74-CU fluorescence microscope.

### PAS (Periodic acid–Schiff) staining

Distal small intestine tissues sections were stained using the Periodic Acid-Schiff (PAS) technique to visualize carbohydrate-rich components. Formalin-fixed, paraffin-embedded tissue sections were deparaffinized in xylene, rehydrated through graded alcohols, and rinsed in distilled water. The sections were then oxidized with 1% periodic acid for 10 minutes, followed by staining with Schiff’s reagent for 15 minutes. After washing in running tap water for 10 minutes, the sections were counterstained with hematoxylin for 2 minutes to visualize nuclei. Finally, the sections were dehydrated through graded alcohols, cleared in xylene, and mounted with a permanent medium. PAS-positive structures were identified by their characteristic magenta color. The images were captured using an Olympus DP74-CU microscope.

### TUNEL Assay for Apoptosis Detection

Apoptotic cells were detected using the One-step TUNEL In Situ Apoptosis Kit (Red, Elab Fluor® 594) (Catalog#E-CK-A322, Elabscience) following the manufacturer’s protocol. Briefly, formalin-fixed, paraffin-embedded distal small intestine tissue sections were deparaffinized, rehydrated through graded alcohols, and permeabilized with Proteinase K (20 μg/mL) for 20 minutes at room temperature. Samples were first incubated with 100 µL of TdT Equilibration Buffer at 37°C for 30 minutes. The equilibration buffer was then removed using absorbent paper, and 50 µL of Labeling Working Solution was added to each slide. Slides were incubated at 37°C for 60 minutes in a humidified chamber, protected from light. After labeling, the slides were washed three times with PBS for 5 minutes each and the sections were counterstained with DAPI to visualize nuclei. A DNase-treated section served as positive control, while negative control was prepared by omitting the TdT enzyme. Slides were mounted using anti-fade mounting medium, and TUNEL-positive cells were visualized via fluorescent microscope (Olympus, model DP74-CU).

### Tamoxifen treatment

6-week-old Rosa26-cre^ERT2^ NLRC4 KI^fl/fl^ mice were treated with tamoxifen (MedChemExpress, HY-13757A, 75 mg/kg body weight) consecutively for 5 days by intraperitoneal injection to obtain global conditional NLRC4 KI mice (Referred to as NLRC4 cKI), Rosa26-cre^ERT2^ NLRC4 KI^fl/fl^ mice treated with corn oil (vehicle) were used as controls.

### Salmonella cultures and infection in vitro

Salmonella Typhimurium strain (14028; ATCC) was grown in LB broth at 37°C for overnight with shaking until mid-log phase (OD600 is about 0.6). Overnight-cultured bacteria were diluted at 1:100 and cultured for another 3 h to induce SPI-1 expression. Bacterial cells were harvested by centrifugation at 4,000 g for 10 min, washed twice with phosphate-buffered saline (PBS), and resuspended to the desired concentration. Bacteria were added to BMDMs at a multiplicity of infection (MOI) of 10 – 50 for 0.5-1 hour.

### Inflammasome activation

To induce inflammasome activation, 2 × 10^6^ BMDM cells were plated in a 6-well plate overnight, and the medium was changed to Opti-MEM the next morning before stimulation. To induce canonical NLRC4 inflammasome activation with two signals (Signal 1 and Signal 2), the cells were first primed with 500 ng/ml LPS (Escherichia coli serotype 0111:B4, Sigma-Aldrich) for 3 h. After that, cells were infected with S. Typhimurium (10 MOI). To induce spontaneous inflammasome activation in the absence of signal 2, the cells were primed with LPS (10 ng/ml) for 16 h. In some indicated experiments, BMDM cells were treated with LRRK2 inhibitors, including LRRK2-IN-1 (Catalog#S7584, Selleckchem) and MLi-2 (Catalog#HY-100411, MedChemExpress), at concentrations of 10 nM. For all experiments, cells were lysed with RIPK buffer supplemented with complete protease inhibitor cocktail (Roche), and cell-free supernatant was either collected for ELISA analysis or concentrated with methanol and chloroform for immunoblot as previously described^21^.

### LDH-Cytotoxicity assay

Lactate dehydrogenase (LDH) activity was measured using LDH-Cytotoxicity Colorimetric Assay Kit (Catalog # K311-400, BioVision) following the manufacturer’s instruction. Briefly, cells were seeded in a 96-well plate at a density of 4 × 10^4^ cells per well and incubated at 37°C. Following the designated treatments, 100 µL of cell culture supernatant was collected and mixed with 100 µL of the LDH assay reagent. The mixture was incubated for 30 minutes at room temperature, protected from light. Absorbance was measured at 490 nm with a reference wavelength of 600 nm using a microplate reader. Maximum LDH release was determined by lysing control cells with 1% Triton X-100 (high control), and Minimums LDH release was determined by medium only wells (low control), the percentage of LDH release was calculated using the following formula: cytotoxicity (%) = (Sample OD495-low control)/(high control-low control).

### Statistic

P values for two group comparisons were determined by Student’s t test. P values for body weight change and survive rate were determined by two-way ANOVA and Log-rank (Mantel-Cox) test, respectively. All values will be presented as means ± SEM, and p values<0.05 will be considered significant.

## Results

### NLRC4 V341A KI mice manifest inflammasome hyperactivation

To generate the NLRC4 V341A knock-in (KI) mice, we replaced the coding sequence (CDS) of exon 2 with the “loxP-stop-loxP kozak-mutant Nlrc4 CDS-3*Flag-IRES-EGFP-polyA” in the target vector, introducing the V341A mutation (GTG to GCG) into the mutant Nlrc4 CDS (**Fig. 1A**). To achieve global expression of the KI alleles, we utilized E2a-Cre transgenic mice, which direct Cre expression to early mouse embryos^16^. E2a^Cre/Cre^ NLRC4 KI^fl/+^ mice were then bred with each other to obtain E2a-Cre NLRC4 KI^fl/fl^ (referred to as NLRC4^KI/KI^ or KI mice) and E2a-Cre NLRC4^+/+^ (wild-type control) littermate controls. The birth of NLRC4 KI mice followed Mendelian inheritance patterns (**Fig. 1B**).

NLRC4 KI and wild-type littermate control mice were genotyped via immunoblotting using antibodies against NLRC4 and Flag Tag. Notably, the expression of NLRC4 KI was comparable to that of endogenous NLRC4 in the colon tissue of wild-type controls (**Fig. 1C**). To further validate NLRC4 V341A KI expression, we analyzed KI-Flag expression in small intestine tissues, intestinal epithelial cells (IECs), and bone marrow-derived macrophages (BMDMs). We observed NLRC4 V341A KI expression across all tested tissues from KI mice, but not in controls, with notably higher NLRC4 expression in IECs compared to BMDMs (**Fig. 1D**).

Given that hyperactivation of the NLRC4 inflammasome has been reported in AIFEC patients^7, 8^, we investigated NLRC4 inflammasome activation in KI mice. IECs from KI mice exhibited significant caspase-1 and GSDMD cleavage, which were absent in wild-type controls (**Fig. 1E**), indicating hyperactivation of the inflammasome in NLRC4 KI mice. Moreover, ELISA analysis revealed significantly elevated levels of IL-1β and IL-18 in both sera and colon explant cultures from KI mice compared to wild-type littermate controls (**Fig. 1F-G**). Collectively, these data suggest that NLRC4 KI mice exhibit spontaneous hyperactivation of the NLRC4 inflammasome.

### NLRC4 V341A KI mice develop autoinflammation

Autoinflammation refers to a group of disorders characterized by uncontrolled inflammation that arises from dysregulation of the innate immune system, rather than from infections, autoimmune processes, or other external triggers^22, 23^. Autoinflammation is a hallmark of AIFEC associated with the NLRC4 V341A mutation in humans^7, 8, 13^. Therefore, we characterized the autoinflammatory phenotype in NLRC4 knock-in (KI) mice. Notably, NLRC4 KI mice exhibited retarded growth (**Fig. 2A-B**), with most succumbing within 10 days postnatally (**Fig. 2B**). Serum ELISA analysis revealed significantly elevated levels of ferritin and IL-6, alongside decreased hemoglobin levels in KI mice (**Fig. 2C**), in addition to heightened IL-1β and IL-18 levels (**Fig. 1G**), indicating systemic autoinflammation. Further blood biochemical analysis indicated multi-organ damage in 6-day-old KI pups, evidenced by elevated AST and ALT levels reflecting liver damage, increased BUN and uric acid levels suggesting kidney impairment, and elevated creatine kinase levels indicative of heart damage or muscle injury (**Fig. 2D**). In contrast, blood glucose levels were markedly reduced in NLRC4 KI mice, reflecting hypoglycemia. However, total protein and albumin levels in the blood of KI mice were higher than those in wild-type (WT) mice, suggesting that hypoglycemia in KI mice was not due to under-nourishment (**Fig. 2D**).

**Figure 2.**
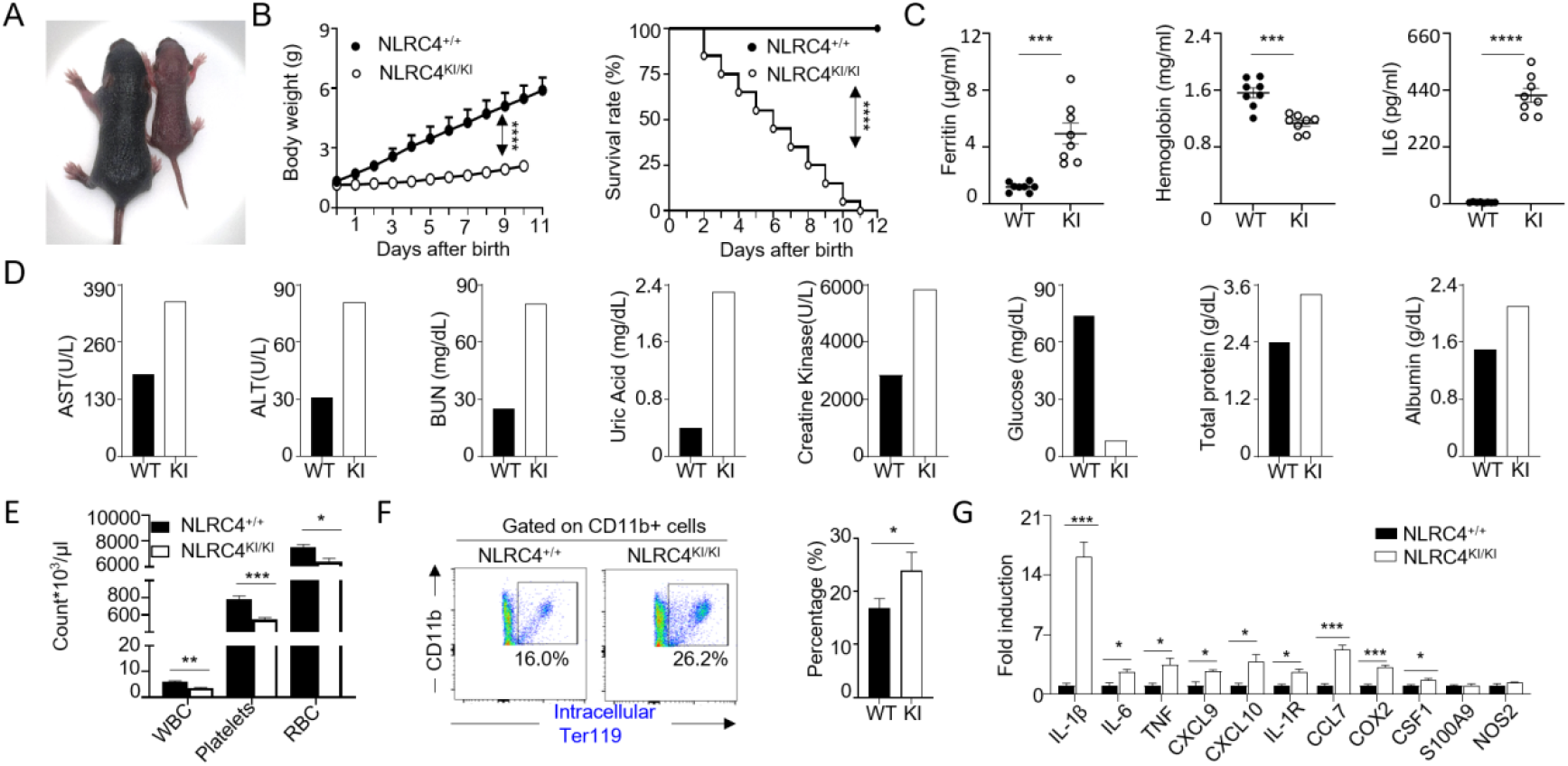
NLRC4 V341A KI mice develop autoinflammation. (A) Gross phenotype of NLRC4 WT (left) and NLRC4 KI (right) mice on day 6 after birth. (B) Body weight (left) and survival rate (right) of NLRC4 WT (NLRC4^+/+^) and NLRC4 KI (NLRC4^KI/KI^) mice after birth were plotted (n=20 per group). P values were determined by two-way ANOVA and Mantel-Cox test, respectively. (C) Levels of serum ferritin, hemoglobin and IL-6 from 6-day-old NLRC4 WT and NLRC4 KI mice were measured by ELISA (n=8 per group). (D) Chemical parameters of blood from NLRC4 WT and KI mice were measured as indicated. ALT, alanine aminotransferase; BUN, blood urea nitrogen; LDH, Lactate Dehydrogenase. Sera were pooled from 8-10 mice per group at the age of 6-day old. (E) Blood cell from 6-day-old NLRC4 WT and NLRC4 KI mice were counted for the indicated cellular parameters (n=5 per group). WBC, white blood cell; RBC, Red blood cell. (F) Hemophagocytosis in spleen was analyzed by flow cytometry. Splenocytes from 6-day-old NLRC4 WT and NLRC4 KI mice were first stained with Fluorescence-conjugated CD11b (myeloid cells); then stained cells were incubated with unconjugated Ter119 antibodies to block corresponding surface antigens; Lastly, cells were permeabilized for intracellular staining of phagocytosed Ter119+ cells (n=3 per group). (G) Isolated macrophages from the spleen of 6-day-old NLRC4 WT and KI pups were analyzed for inflammatory gene expression as indicated by Q-PCR (n=4 per group). p values were determined by Student’s t test, *p<0.05, **p<0.01, ***p<0.001 and **** p<0.0001. Data are shown as mean ± SEM. Data are representative of three independent experiments (panels C, E, F).

The autoinflammation observed in AIFEC is characteristic of Macrophage Activation Syndrome (MAS), which often manifests as cytopenia, macrophage hyperactivation, and hemophagocytosis ^7, 8^. Consistent with clinical observations, we found decreased leukocyte, red blood cell, and platelet counts in the blood of NLRC4 KI mice (**Fig. 2E**). Intracellular staining of splenocytes followed by flow cytometric analysis revealed evidence of hemophagocytosis, as indicated by a significant increase in the frequency of Ter119+ cells within CD11b+ myeloid cells in KI mice compared to WT controls (**Fig. 2F**). Additionally, Q-PCR analysis of sorted macrophages from the spleens of NLRC4 KI and WT mice demonstrated that macrophages from KI mice were hyperactivated, producing significantly higher levels of pro-inflammatory cytokines and chemokines (**Fig. 2G**). Collectively, these findings suggest that NLRC4 KI mice develop autoinflammation.

### NLRC4 V341A KI mice develop infantile enterocolitis

A hallmark of AIFEC is infantile enterocolitis. Patients with AIFEC often present diarrhea, abdominal pain, villus blunting, and mixed inflammatory infiltrates in the intestine^8^. To determine whether NLRC4 KI pups develop infantile enterocolitis, we collected intestinal tissues from 6-day-old NLRC4 KI and wild-type (WT) control mice. H&E staining of the distal small intestine and colon revealed villous blunting and epithelial damage, along with increased inflammatory cell infiltration in the lamina propria of KI mice (**Fig. 3A**). Flow cytometric analysis of mononuclear cells isolated from colon lamina propria showed an increase in macrophages and neutrophils, accompanied by a decrease in T cells in the KI mice (**Fig. 3B**). Furthermore, Q-PCR analysis of colon tissue from KI mice indicated elevated expression of pro-inflammatory cytokines and chemokines compared to WT controls (**Fig. 3C**). ELISA assays of supernatants from colon explant cultures demonstrated significantly higher levels of IL-6 and TNF-α in KI mice, in addition to increased IL-1β and IL-18 (**Fig. 1F**). Notably, NLRC4 KI mice exhibited enlarged colons and increased colon weight (**Fig. 3E; Fig. 3F**, left two panels). These mice also showed significant diarrhea, as evidenced by a higher wet/dry feces ratio and stool score, with a higher score indicating more severe diarrhea (**Fig. 3F**, right two panels). Taken together, these findings suggest that NLRC4 KI mice develop infantile enterocolitis, mirroring the gastrointestinal symptoms observed in AIFEC patients.

**Figure 3.**
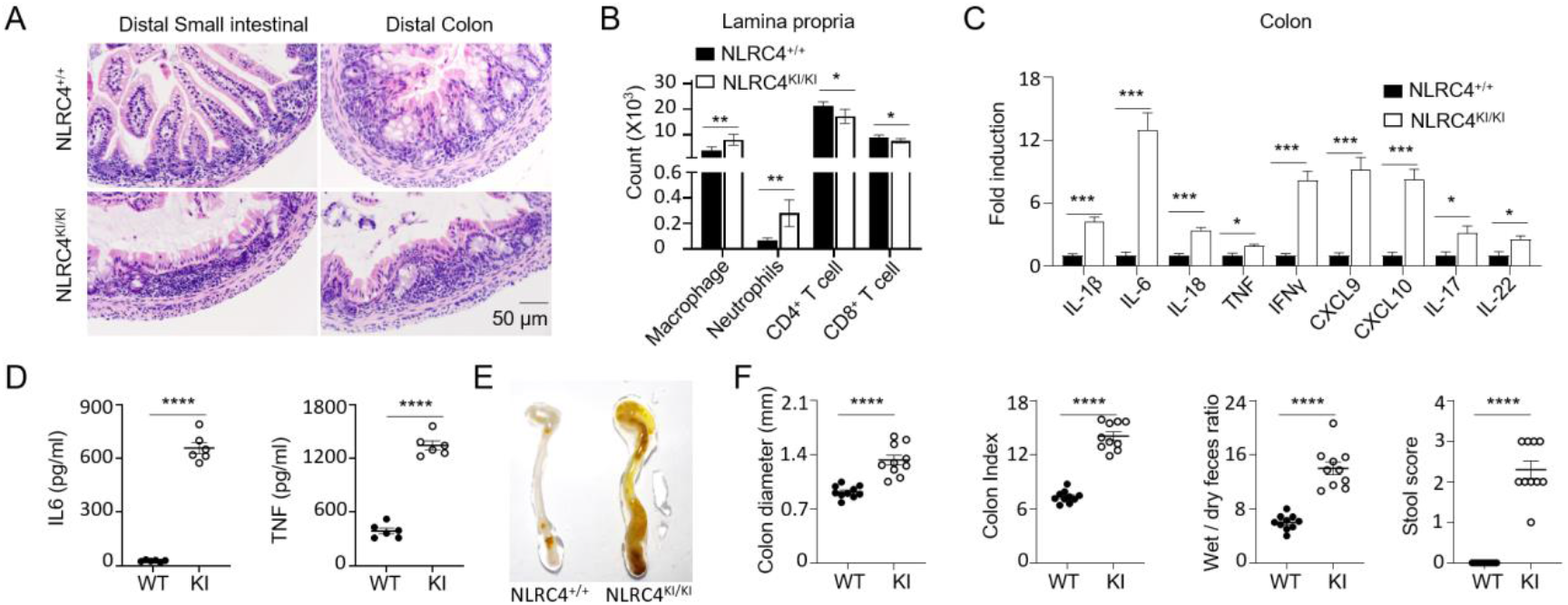
NLRC4 V341A KI mice develop infantile enterocolitis. (A) Representative H&E staining of small intestine and colon tissues from 6-day-old NLRC4 WT and KI mice as indicated. Inflammatory cell infiltration in lamina propria from colon tissues of 6-day-old NLRC4 WT and KI mice were analyzed by flow cytometry (n=5 per group). (C) Colon tissues from 6-day-old NLRC4 WT and KI mice were analyzed for inflammatory gene expression as indicated by real-time PCR (n=4 per group). (D) IL-6 and TNF levels in supernatant of colon explant culture from 6-day-old NLRC4 WT and KI mice were measured by ELISA (n=6 per group). (E) Representative images of gross colon from 6-day-old NLRC4 WT and KI mice. (F) Colon diameter, Colon index (colon weight/body weight), Wet/dry feces ratio and Stool score of 6-day-old NLRC4 WT and KI mice are shown as indicated (n=10 per group). Data are shown as mean ± SEM. p values were determined by Student’s t test, *p<0.05, **p<0.01, ***p<0.001 and **** p<0.0001. Data are representative of three independent experiments.

### Gut barrier integrity is impaired in NLRC4 V341A KI mice

Excessive IL-18 production, as seen in conditions involving deletion of IL-18BP or colitis, has been linked to the loss of mucus-producing goblet cells^24^. Given that NLRC4 KI mice exhibit overproduction of IL-18 in the gut, we assessed goblet cell populations in the small intestine using PAS staining. This analysis revealed a dramatic reduction of goblet cells in the KI tissue compared to WT controls. Interestingly, we also observed a decrease in Paneth cells in the small intestines of KI mice (**Fig. 4A**, left two panels). As shown in **Fig. 1E**, intestinal epithelial cells (IECs) from KI tissue exhibited robust GSDMD cleavage, which is typically associated with pyroptosis. To investigate cell death in the gut tissue, we performed a TUNEL assay, revealing significant cell death in the KI tissue while barely any was observed in the WT controls (**Fig. 4A**, panel 3).

**Figure 4.**
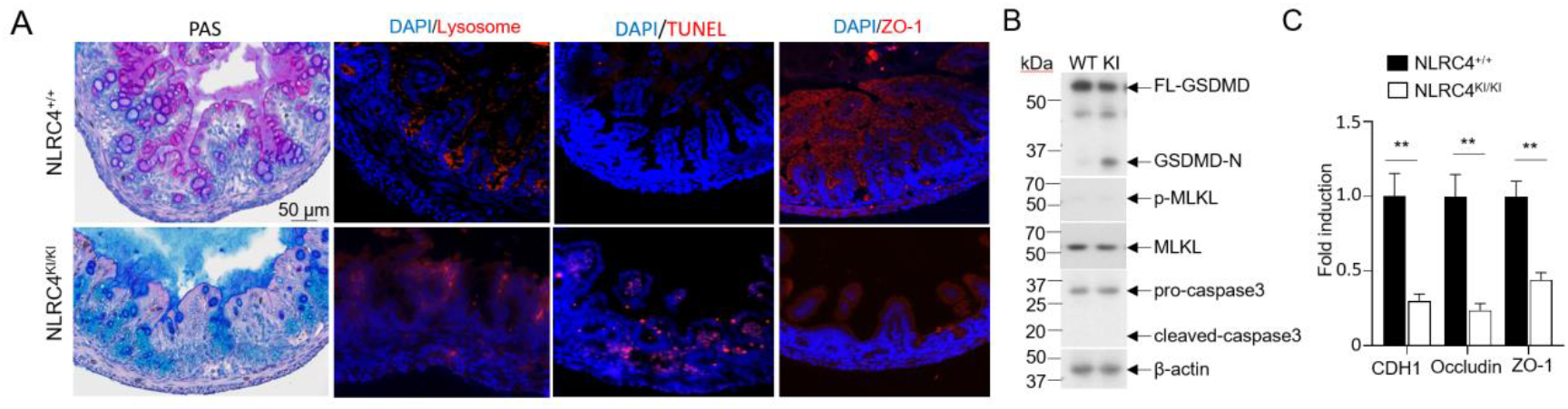
Gut barrier integrity is impaired in NLRC4 V341A KI mice. (A) Representative images for different histochemical staining as indicated, samples were distal small intestine tissues from 6-day-old NLRC4 WT and NLRC4 KI mice. PAS (Periodic acid–Schiff) staining for goblet cells, lysosome for Paneth cells, TUNEL for cell death and ZO-1 for tight junction assessment. (B) Western blot determining cell death type of distal small intestine tissues from 6-day-old NLRC4 WT and NLRC4 KI mice. Cleaved Gasdermin D (GSDMD-N) is indicative of pyroptosis, p-MLKL is a marker of necroptosis, cleaved caspase 3 signify apoptotic cell death. Representative tight junction genes as indicated were analyzed by Q-PCR. Samples were intestinal epithelial cells isolated from 6-day-old NLRC4 WT and NLRC4 KI mice (n=4 per group). p values were determined by Student’s t test, **p<0.01. Data are shown as mean ± SEM. Data are representative of three independent experiments.

Additionally, Western blot analysis of IECs from both WT and KI mice indicated no differences in necroptosis (assessed by probing for pMLKL) or apoptosis (assessed by probing for cleaved caspase-3) between the two groups (**Fig. 4B**), suggesting that the increased cell death in the gut epithelium of KI mice is likely due to pyroptosis. Inflammasome activation can also modulate the expression of tight junction proteins via IL-1β or IL-18^25, 26^. We analyzed tight junction proteins in the small intestines of WT and KI mice using immunofluorescent staining for ZO-1, which showed a dramatic reduction in ZO-1 levels in the KI mice compared to WT controls (**Fig. 4A**, panel 4). Q-PCR analysis for CDH1 (cadherin), occludin and ZO-1 further confirmed that the expression levels of these tight junction proteins were significantly decreased in the intestinal epithelial cells of KI mice relative to those in WT controls (**Fig. 4C**). Collectively, these findings suggest that the integrity of the gut barrier is impaired in NLRC4 KI mice.

### NLRC4 V341A KI mice develop AIFEC after birth

While we observed severe infantile enterocolitis and autoinflammation in NLRC4 KI mice at 6 days postnatally (**Fig. 1-4**), it remained unclear whether these mice developed disease earlier, before birth. To investigate this, we monitored disease progression at postnatal days 0, 3, and 6, using littermate wild-type (WT) controls at each time point. Initially, H&E staining of intestinal tissues revealed no significant damage in either the small intestine or colon of NLRC4 KI mice at birth. However, by day 3, we observed crypt disruption and inflammatory infiltrates in the intestines of NLRC4 KI mice, which worsened by day 6 (**Fig. 5A**).

**Figure 5.**
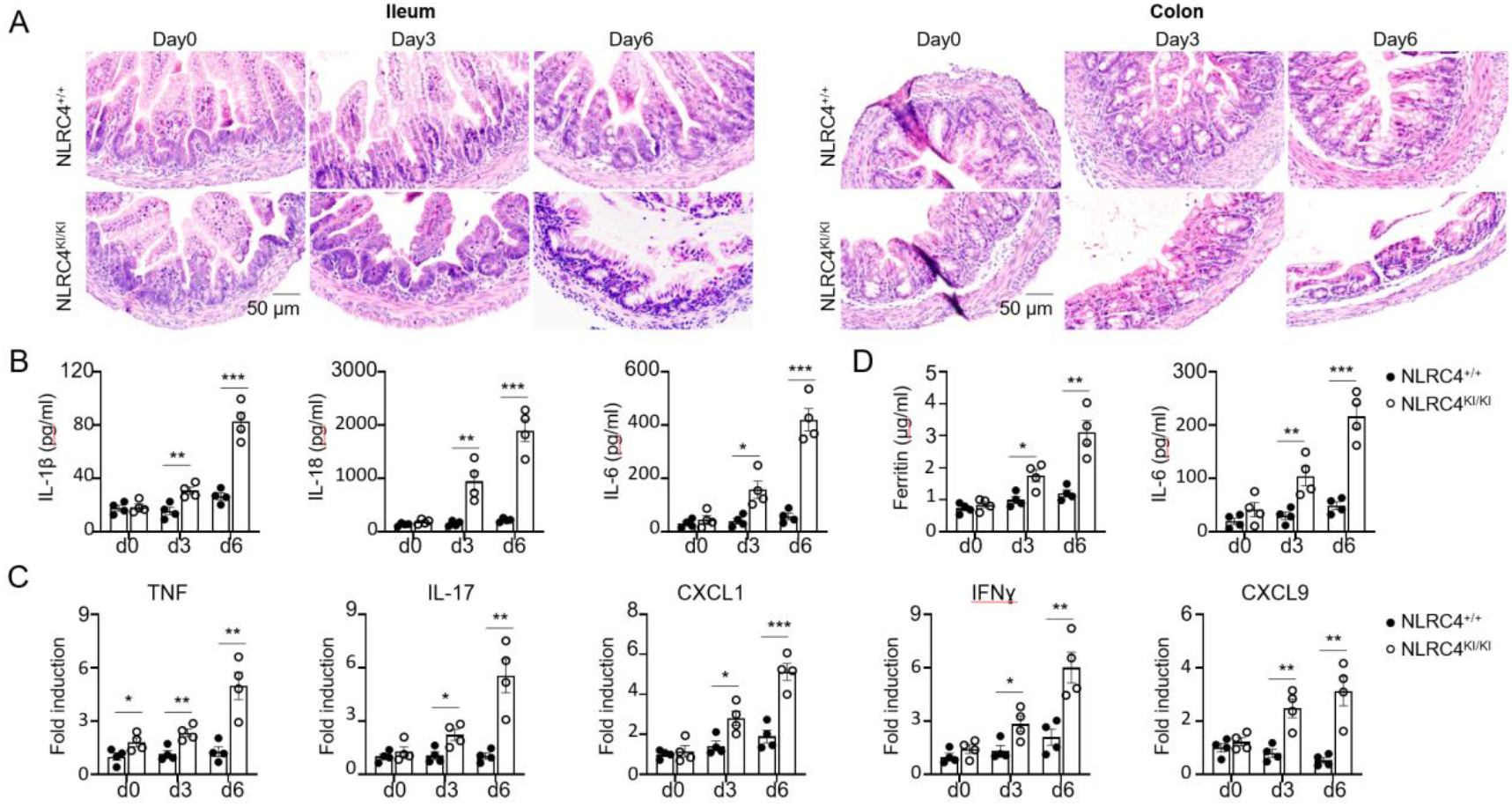
Kinetic analysis of AIFEC development in NLRC4 V341A KI pups. (A) Representative H&E staining of small intestine (left) and colon tissues (right) from 0,3,6-day-old NLRC4 WT and NLRC4 KI mice. (B) IL-1β, IL-18 and IL-6 levels from supernatant of colon explant culture of NLRC4 WT and KI mice were measured by ELISA (n=4 per group). (C) Colon tissues from 0,3,6-day-old NLRC4 WT and NLRC4 KI mice were analyzed for inflammatory gene expression as indicated by Q-PCR (n=4 per group). (D) Levels of serum ferritin and IL-6 from 0,3,6-day-old NLRC4 WT and NLRC4 KI mice were measured by ELISA (n=4 per group). Data are shown as mean ± SEM. p values were determined by Student’s t test, *p<0.05, **p<0.01 and ***p<0.001.

Consistent with these histological findings, ELISA analysis of supernatants from colon explant cultures showed no differences in IL-1β, IL-18, and IL-6 levels between the two groups at day 0. However, significant increases were noted in NLRC4 V341A pups at postnatal day 3 and 6 (**Fig. 5B**). This was further supported by qPCR analysis, which revealed elevated expression of pro-inflammatory cytokines (TNF, IL-17, and IFN-γ) and chemokines (CXCL1 and CXCL9) in KI pups (**Fig. 5C**). Additionally, we analyzed serum ferritin and IL-6 levels to assess potential autoinflammation in early infancy. Both serum ferritin and IL-6 levels were comparable between NLRC4 V341A and WT pups at day 0, but significantly elevated in the KI group as the disease progressed at postnatal days 3 and 6 (**Fig. 5D**), indicating a progression of systemic autoinflammation. Collectively, these data suggest that AIFEC in NLRC4 KI mice develops after birth rather than before.

### Adult NLRC4 V341A conditional KI mice exhibit autoinflammation with mild colitis

Infant AIFEC patients exhibit severe enterocolitis. In contrast, colitis in adult patients typically resolves or becomes much milder, though autoinflammation persists into adulthood^13, 27^. Constitutive NLRC4 KI pups, however, die within 10 days postnatally, preventing further study of autoinflammation in adult KI mice. To model the adult AIFEC patient phenotype, we employed R26-Cre^ER^ mice^28^ to induce global NLRC4 KI expression in adult mice using tamoxifen. To test whether adult NLRC4 conditional KI mice developed autoinflammation and enterocolitis, 6-week-old R26-Cre^ER^ NLRC4 V341A^fl/fl^ mice were treated with tamoxifen for 5 consecutive days to generate conditional NLRC4 KI mice (NLRC4 cKI). Littermate controls were R26-Cre^ER^ NLRC4 V341A^fl/fl^ mice treated with corn oil (vehicle). Mouse body weight and survival were monitored daily following tamoxifen treatment. The NLRC4 cKI mice exhibited normal survival rates and showed no significant changes in body weight compared to the control group (data not shown). At 8 weeks of age, both NLRC4 cKI and control mice were euthanized for genotyping and phenotyping. Western blot analysis of small intestine tissue and IECs confirmed NLRC4 KI expression in the NLRC4 cKI mice (**Fig. 6A**).

**Figure 6.**
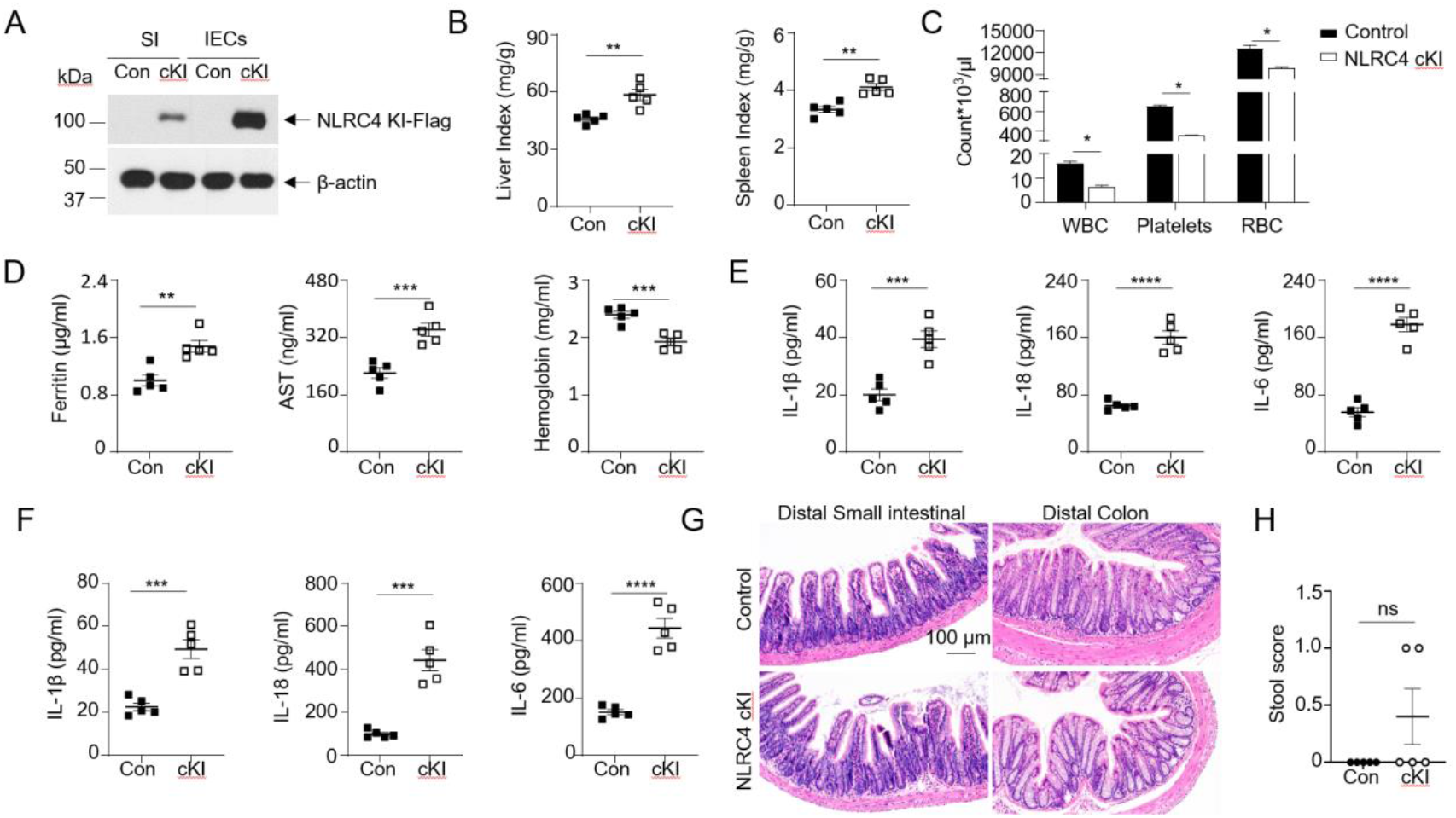
Adult NLRC4 cKI mice exhibit autoinflammation with mild colitis. 6-weeks-old Rosa26-cre^ER^ NLRC4 V341A^fl/fl^ mice were treated with tamoxifen (75 mg/kg) consecutively for 5 days to obtain global conditional NLRC4 KI (NLRC4 cKI) mice, the mice treated with corn oil (vehicle) were used as littermate controls. After treatment, mice were sacrificed at 8 weeks old for the following analysis. (A) NLRC4 cKI and control mice were genotyped by western blot. Samples from small intestines (SI), Intestinal epithelial cells (IECs) were probed by Flag and β-actin antibodies as indicated. (B) Liver index (liver weight/body weight, Left) and spleen index (Right) of NLRC4 cKI and control mice are presented (n=5 per group). (C) Blood cells from 8-week-old NLRC4 cKI and control mice were analyzed by flow cytometry as indicated (n=5 per group). WBC, white blood cell; RBC, Red blood cell. (D) Serum Ferritin, AST and hemoglobin levels of NLRC4 cKI and control mice were measured by ELISA as indicated (n=5 per group). (E) Serum IL-1β, IL-18 and IL-6 levels of NLRC4 cKI and control mice were measured by ELISA as indicated (n=5 per group). (F) IL-1β, IL-18 and IL-6 levels in supernatants of colon explant culture from NLRC4 cKI and control mice were measured by ELISA as indicated (n=5 per group). (G) Representative H&E staining of small intestine and colon tissues from 8-week-old NLRC4 cKI and control mice as indicated. (H) Stool scores of 8-weeks-old NLRC4 cKI and control mice. Data are shown as mean ± SEM. p values were determined by Student’s t test, *p<0.05, **p<0.01, ***p<0.001 and **** p<0.0001. Data are representative of at least two independent experiments.

Phenotypically, NLRC4 cKI mice exhibited splenomegaly and hepatomegaly (**Fig. 6B**). Additionally, blood analysis revealed cytopenia, indicated by reduced blood cell counts (**Fig. 6C**). Serum levels of ferritin, AST, IL-1β, IL-18, and IL-6 were significantly elevated in NLRC4 cKI mice compared to controls (**Fig. 6D-E**), while hemoglobin levels were markedly reduced, further supporting the presence of robust autoinflammation in these mice (**Fig. 6D**). To assess intestinal inflammation, colon explant cultures were performed, revealing a significant increase in IL-1β, IL-18, and IL-6 levels in the supernatants of NLRC4 cKI mice compared to vehicle-treated controls (**Fig. 6F**). Real-time PCR also showed upregulated expression of inflammatory cytokines and chemokines in colon tissues from NLRC4 cKI mice (data not shown). However, despite these inflammatory markers, we did not observe substantial damage or inflammatory infiltrates in the gut epithelium of adult NLRC4 cKI mice, which contrasts with the severe epithelial damage seen in infantile NLRC4 KI mice (**Fig. 6G**). Additionally, no significant differences were observed in stool scores between NLRC4 cKI mice and WT controls (**Fig. 6H**). In summary, these findings suggest that adult NLRC4 cKI mice exhibit autoinflammation with mild colitis, serving as a relevant model for adult AIFEC patients.

### LRRK2 kinase inhibitors prevent excessive NLRC4 V341A inflammasome activation

Previous studies have demonstrated that gain-of-function mutations in NLRC4 lead to hyperactivation of the NLRC4 inflammasome in macrophages derived from AIFEC patients^7, 8^. We hypothesized that a similar mechanism occurs in AIFEC mice. To test this, we first induced canonical NLRC4 inflammasome activation in bone marrow-derived macrophages (BMDMs) from WT and NLRC4 KI mice using two signals (Signal 1: LPS priming; Signal 2: Salmonella infection). The V341A mutation significantly enhanced inflammasome activation in this context, as shown by markedly elevated levels of cleaved IL-1β, Caspase-1, and GSDMD (**Fig. 7A**). Previously, we demonstrated that LRRK2 kinase activity is critical for NLRC4 inflammasome activation ^29^. Building on this, we investigated whether inhibiting LRRK2 kinase activity could mitigate the hyperactivation of the NLRC4 V341A inflammasome. Strikingly, the excessive inflammasome activation triggered by the two-signal stimulation was effectively suppressed by the LRRK2 kinase inhibitors LRRK2-IN-1 and MLi-2 (**Fig. 7B**).

**Fig.7.**
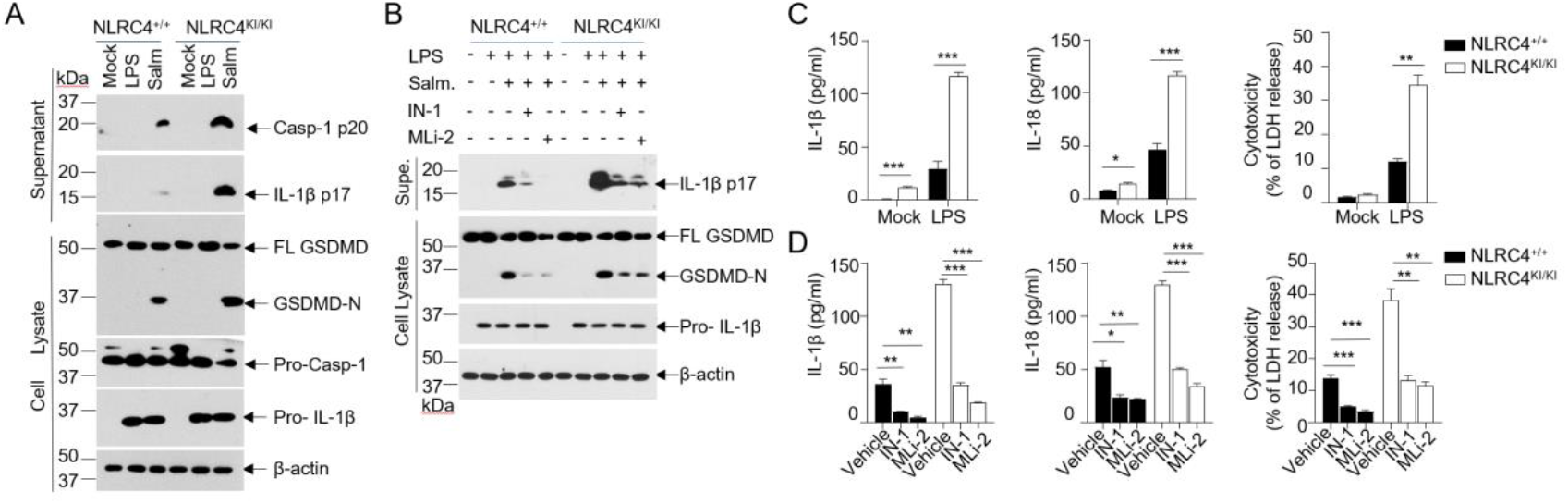
LRRK2 kinase activity is crucial for NLRC4 V341A inflammasome hyperactivation. BMDMs were cultured from adult R26Cre^ER^ NLRC4^fl/fl^ mice after tamoxifen induction in vivo, BMDMs from WT control mice were included as control. (A) Western blot analysis of inflammasome activation in bone marrow-derived macrophages (BMDMs) from NLRC4 wild-type (WT) and knock-in (KI) mice. BMDMs were primed with 500 ng/ml LPS for 3 hours, followed by infection with Salmonella Typhimurium (Salm) at a multiplicity of infection (MOI) of 10 for 30 minutes. Cell lysates and culture supernatants were immunoblotted with the indicated antibodies. (B) Western blot analysis of the role of LRRK2 kinase in inflammasome activation in both NLRC4 WT and KI BMDMs. Cells were treated with LRRK2 kinase inhibitors (LRRK2-IN-1 or MLi-2). BMDMs were primed for 3 hours in media containing 500 ng/ml LPS, treated with 5 μM LRRK2-IN-1 or MLi-2 for 2 hours, and then infected with S. Typhimurium at an MOI of 10 for 30 minutes, followed by western blot analysis. (C) ELISA analysis of constitutive inflammasome activation in BMDMs from NLRC4 WT and KI mice. BMDMs were primed with 10 ng/ml LPS for 18 hours. (D) Role of LRRK2 kinase activity in constitutive NLRC4 V341A inflammasome activation. NLRC4 WT and KI BMDMs were primed with 10 ng/ml LPS for 18 hours and concurrently treated with 5 μM LRRK2-IN-1 or MLi-2 as indicated. Supernatants were collected and analyzed by ELISA. Lactate dehydrogenase (LDH) release is reported relative to levels induced by complete cell lysis with Triton X-100 (n=3/group). Data are shown as mean ± SEM. p-values were determined using Student’s t-test (*p < 0.05, **p < 0.01, ***p < 0.001). Data are representative of three independent experiments.

Furthermore, monocyte-derived macrophages from AIFEC patients with the NLRC4 V341A mutation exhibit constitutive inflammasome activation following low-dose LPS exposure for 18 hours ^8^. Consistently, we observed constitutive inflammasome activation in BMDMs from NLRC4 V341A mice, but not in WT controls, as indicated by the substantial increase in IL-1β, IL-18, and lactate dehydrogenase (LDH) production (**Fig. 7C**). Notably, this spontaneous inflammasome activation was also abrogated by LRRK2 kinase inhibitors LRRK2-IN-1 and MLi-2 (**Fig. 7D**). In conclusion, LRRK2 kinase inhibitors effectively attenuate both spontaneous and canonical NLRC4 V341A inflammasome activation, highlighting the therapeutic potential of targeting LRRK2 kinase activity for the treatment of autoinflammation in infantile enterocolitis.

## Discussion

While NLRC4 inflammasome activation is critical for host defense, excessive NLRC4 inflammasome activation can lead to severe diseases, which is evidenced by decreased survival, severe intestinal bleeding and multiple organ damage induced by E.coli O21: H^+^-induced NLRC4 inflammasome overactivation^30^. This phenotype was recapitulated by systemic activation of NLRC4 inflammasome via injecting mice with bacterial needle/rod proteins^31^. Furthermore, eicosanoids produced upon NLRC4 inflammasome activation have been shown to contribute to enterocolitis^31^. These seminar studies suggested that unchecked NLRC4 inflammasome activation may be detrimental in animal models. However, direct evidence linking excessive NLRC4 inflammasome activation to human disease comes from two pioneering reports in 2014, which identified gain-of-function mutations in NLRC4 (specifically T337S and V341A) as contributors to the pathogenesis of autoinflammatory enterocolitis (AIFEC) ^7, 8^. Since then, the spectrum of associated mutations has expanded ^9-12^.

Previous attempts to establish animal models for AIFEC include the generation of an NLRC4 T337S knock-in (KI) mouse, which, despite elevated IL-18 levels, does not spontaneously develop disease^32^. Consequently, the lack of a spontaneous model has limited our understanding of AIFEC pathogenesis and the development of effective treatments. In this study, we present a conditional NLRC4 V341 KI mouse model, where pups spontaneously develop severe infantile enterocolitis and autoinflammation. In contrast, adult conditional KI mice predominantly exhibit autoinflammation with only mild enterocolitis. AIFEC is characterized by life-threatening enterocolitis during infancy and recurrent, severe autoinflammation throughout life ^5, 13^. Our NLRC4 KI model effectively recapitulates these key features, with infant mice succumbing within 10 days post-birth, closely mirroring the high mortality rates seen in infant AIFEC patients, many of whom do not survive beyond early childhood without appropriate medical intervention. This underscores the significance of our novel animal model for studying AIFEC. Interestingly, a recent report by Wullaert et al. described an NLRC4 V341A KI mouse that did not develop the AIFEC phenotype, though elevated IL-18 levels were noted ^33^. The reason for this discrepancy remains unclear but may be attributed to differences in the mouse models. The Wullaert group utilized Crispr-Cas9 technology^33^ which may increase the concern about off-target effect^34-36^, while our model was generated using Cyagen’s TurboKnockout® gene-targeting service with traditional homologous recombination. Additionally, in the Wullaert report, NLRC4 KI protein expression was not confirmed by western blot or other methods^33^, leaving uncertainty about NLRC4 KI protein expression. In contrast, we validated NLRC4 KI expression at the protein level using both a Flag Tag antibody and an endogenous NLRC4 antibody. Furthermore, environmental factors, such as differences in animal housing conditions, could also influence phenotypic outcomes.

Our model not only serves as a novel system for studying AIFEC but also has substantial potential for investigating very early-onset inflammatory bowel disease (VEO-IBD). VEO-IBD, diagnosed in individuals under the age of six, is a heterogeneous and severe disorder with potentially fatal outcomes. The incidence of VEO-IBD is rising, but approved therapies remain limited, and traditional treatments for IBD often prove ineffective^37, 38^. Over 75 distinct single-gene defects have been identified as molecular causes of VEO-IBD, accounting for approximately 7.85-7.9% of cases^38-40^. The disease is influenced by a complex interplay of environmental factors, immune dysregulation, impaired gut barrier function, and dysbiosis in genetically susceptible individuals ^41^. Several of these monogenic defects have been associated with dysregulated inflammasome activity, underscoring the central role of inflammasomes in maintaining intestinal homeostasis ^42^. The discovery of de novo gain-of-function mutations in NLRC4, which lead to AIFEC, established a direct connection between IBD pathogenesis and dysregulated inflammasome activation^8, 40, 42, 43^. However, animal models of monogenic VEO-IBD driven by excessive inflammasome activation remain limited. AIFEC patients typically present with infantile-onset IBD before the age of two, with symptoms including diarrhea, abdominal pain, villus blunting, and mixed inflammatory infiltrates in both the small intestine and colon^8, 13^. Our NLRC4 KI mice spontaneously develop similar enterocolitis, characterized by gut inflammation, impaired intestinal barrier integrity, disrupted gut epithelium, and severe diarrhea. Therefore, our human-relevant NLRC4 knock-in mouse model of AIFEC offers a unique and valuable tool for studying the pathogenesis of VEO-IBD.

Given that AIFEC is a rare and often severe condition at diagnosis, mechanistic studies and therapeutic explorations in clinical settings are challenging. Our NLRC4 KI model provides a unique opportunity to bridge this gap. With the rising incidence of VEO-IBD, this novel KI model will have broad implications for the study of the pathogenesis and translational research of VEO-IBD, particularly infantile-onset forms of the disease.

## Authors’ Disclosures

The authors declare no conflict of interests.

## Authors’ Contributions

Y.W. designed and performed most of the experiments, interpreted the data, and wrote part of the manuscript. J.G. assisted in mouse genotyping, immunoblotting, and real-time PCR. Y.X. instructed and performed pathology analysis. P.G. and S.S contributed to data interpretation and manuscript revision. Z.K. was integral for experimental design, manuscript writing, data interpretation and project coordination.

## Acknowledgments

This work was supported by Startup Funds from the Department of Pathology, Carver Collage of Medicine, University of Iowa.

## Notes

### Competing Interest Statement

The authors have declared no competing interest.

### Summary of Updates

In Figure 1D, the arrows were pointing incorrectly. I have now corrected them.

